# Alzheimer’s Disease-Related Mutations in APP Influence Colorectal Tumor Formation in a Sex-Dependent Manner

**DOI:** 10.1101/2025.10.31.685949

**Authors:** Takako Ishida-Takaku, Mona Sohrabi, Heidi L. Pecoraro, Colin K. Combs

## Abstract

Cancer and Alzheimer’s disease (AD) are major age-associated diseases. Numerous studies have indicated common mechanisms and biological interactions between these conditions. Furthermore, epidemiological data demonstrate negative correlations between several cancers and AD. Despite this, few studies have explored their mutual influence on pathological outcomes. We utilized a human APP mutant knock-in AD mouse model to investigate how familial mutations influence colorectal cancer development. Using a colitis-associated colorectal cancer model, APP mutations promoted colon tumor formation in male mice but inhibited it in female mice. Correspondingly, inflammatory changes were reduced in the colons of female mice. Transcriptome analysis revealed differential expression, particularly in the enrichment of neuronal marker, steroid hormone, and immune cell signaling pathways. Additionally, distinct macrophage subtypes and neuronal profiles were observed in the colons of male and female APP mutant mice. These findings provide the first elucidation of the sex-dependent effects of APP mutations on colon cancer formation.

## Introduction

Cancer and Alzheimer’s disease (AD) are two of the most prevalent age-associated diseases, presenting significant health challenges worldwide. Epidemiological studies consistently reveal an inverse relationship between these conditions^1–4^. However, data from a large cohort of colorectal cancer patients also suggest that individuals with this type of cancer may have an increased risk of developing AD^5,6^, suggesting intricate biological mechanisms underpin their distinct pathophysiologies. Importantly, sex-dependent differences are well-documented in the epidemiological data for both diseases^7–9^. Hormonal factors appear to be key influencers of disease susceptibility and progression^10^. In colorectal cancer, for instance, the risk is typically lower in women, possibly due to protective effects conferred by higher levels of estrogen, progesterone, or other sex dependent factors^11,12^. The interaction between sex specific factors and AD is particularly evident in the processing of the amyloid precursor protein (APP), which plays a critical role in β-amyloid (Aβ) accumulation^13^, a hallmark of AD pathology. This intersection is crucial as it highlights how sex dependent factors might influence AD progression and, conversely, how AD-related pathways might impact cancer dynamics. Elevated levels of APP and its cleavage product, sAPPα, have been observed in various cancers^14–17^, suggesting that APP might have more than a role in neuronal disorders but also a potential mediator of oncogenic processes. This raises the possibility that APP overexpression or its aberrant processing could contribute to tumorigenic pathways in the colon, particularly through modulation of cellular proliferation and apoptosis^18^. Despite the intriguing associations between APP biology and cancer progression, the underlying mechanisms, particularly those detailing sex-specific differences in disease progression and APP metabolism in the context of colon cancer, remain largely unexplored. This gap in knowledge represents a critical area of research that could uncover potential therapeutic targets for both diseases.

To address this need, our study utilized the *App^NL-G-F^*amyloidosis AD mouse model^19^, which features three familial AD-causing mutations knocked into the mouse App gene. This model provides a unique platform to study the direct effects of human-like amyloid pathology on colorectal cancer development in a controlled genetic background. Our investigations revealed sex-dependent variations in the development of colitis-associated colorectal cancer. Male *App^NL-G-F^* mice displayed an increased tumor burden characterized by a greater number and size of tumors compared to their wild-type counterparts. In contrast, female *App^NL-G-F^* mice exhibited a notable resistance to tumor formation, suggesting a protective effect mediated possibly through sex dependent factors and/or different pathways of APP processing. Gene expression data confirmed male and female specific differences. Genes related to steroid hormone biosynthesis, metabolism, and immune signaling pathways were significantly altered in *App^NL-G-F^* colons compared to those in wild-type mice. Consistent with the gene expression changes, variations in macrophage subtypes and neuron marker expression were also observed in both the colon tumor microenvironments and non-tumorigenic areas. These results suggest that APP influences colon tumorigenesis in a sex dependent manner.

## Results

### Colon tumor formation was inhibited in female *App^NL-G-F^* mice but stimulated in males

We first investigated the expression levels of APP in human colon cancer. Hematoxylin and eosin (H&E) staining was used to compare normal human tissue alongside well-differentiated, moderately differentiated, and poorly differentiated colon tumors in cases of colorectal cancer (Fig. 1A). Immunostaining of the same normal human colon and a range of differentiated tumors showed expression of APP in intestinal epithelial cells (Fig. 1B). These data confirm that APP is expressed in colonic epithelial cells independently of whether the condition is healthy or tumorigenic. Clinical data indicate a distinct correlation between APP expression levels and survival in colorectal cancer patients (Fig. 1C). Specifically, overexpression of APP in male patients is associated with poorer overall survival, while this correlation is weaker in female patients (Fig. 1C). When relapse-free survival was analyzed, female patients with high APP expression showed better outcomes, whereas male patients with high APP expression still exhibited worse relapse-free rates (Fig. S1, A and B). This suggests that APP, and potentially other AD-associated genes, may play a role in tumorigenesis. To investigate the association between CRC and AD, we induced CRC in wild type and *App^NL-G-F^*male and female mice using AOM/DSS treatment (Fig. 1E). The Kaplan-Meier survival curves were used for examining survival rates of wild type and AOM/DSS-treated mice (Fig. 1, F and G). By week 2, the survival rates for the male AOM/DSS-treated groups dropped to 91%, and 83% for wild type, and *App^NL-G-F^* mice, respectively. The survival rate for female wild types decreased from 91% in week 1 to 73% in week 2. In the same week, all the AOM/DSS treated female *App^NL-G-F^* mice survived throughout the study.

**Fig. 1.**
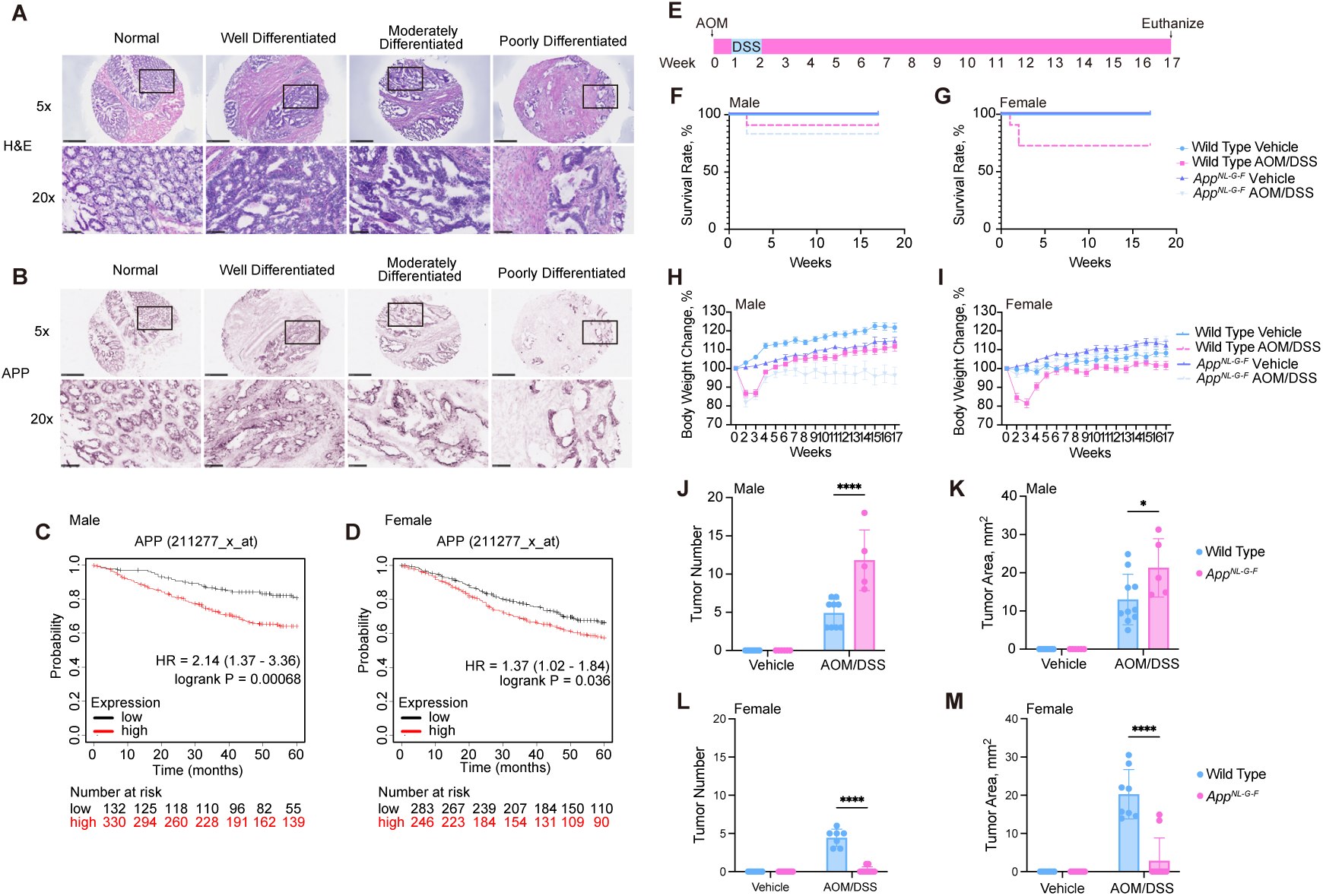
Colon tumor formation was inhibited in female *App^NL-G-F^* mice but stimulated in males. **A**, Histology and APP immunohistochemistry of human normal colon and colorectal cancer tissue arrays. Representative H&E staining of human normal colon and colorectal cancer ranging from well to poorly differentiated tumors are shown with 5X magnification and 20X magnification. **B**, Representative APP immunoreactivity demonstrated APP expression in epithelial cells in human normal colon tissue and colorectal cancer tumors shown with 5X magnification and 20X magnification. **C**,**D** Male (**C**) and female (**D**) Kaplan-Meier Survival Curves for Overall Survival (OS) by APP Expression. **E,** Experimental design and timeline of AOM/DSS treatment. **F**,**G**, The survival rate of male (**F**) and female (**G**) vehicle or AOM/DSS treated wild type and *App^NL-G-F^* mice. **H**, The percentage of body weight changes for male (**H**) and female (**I**). **J**,**K,** Tumor number and area were counted and measured, respectively, from wild type and *App^NL-G-F^*males to indicate the extent of disease induced by AOM/DSS exposure across strain and sex, *p<0.05, ** p<0.01, ***p<0.001, and ****p<0.0001 (mean ± SEM, n=5-11 animals). **L**,**M,** Tumor number and area were counted and measured, respectively, from wild type and *App^NL-G-F^* females to indicate the extent of disease induced by AOM/DSS exposure across strain and sex, *p<0.05, ** p<0.01, ***p<0.001, and ****p<0.0001 (mean ± SEM, n=9-10 animals).

The percentage of weight loss was only graphed for mice that survived the entire procedure (Fig. 1, H and I). AOM/DSS treated groups significantly increased their percentage of weight loss for both males and females in week 2 compared to their respective vehicle controls and started gaining weight from week 4. There were no significant differences in the final body weights of *App^NL-G-F^*males and wild type females in AOM/DSS treated groups from weeks 6 to the end of the study. These results suggest that response to AOM/DSS treatment varies by gender and genotype. The inflammation by DSS exposure is related to increases in colon and spleen weights as well as a decrease in colon length^20^. AOM/DSS treated *App^NL-G-F^* male groups showed a significant increase in colon weights compared to their respective control (Fig. S1, C to E). In females, AOM/DSS treated wild type mice showed shorter colon length and increased colon and spleen weights compared to their *App^NL-G-F^* group (Fig. S1, F to H). These data suggest CAC inflammation affects colon and spleen weights, and colon length in a sex dependent manner.

Next, we counted tumor number and measured tumor area (Fig. 1. J to M). In the male AOM/DSS treated groups, *App^NL-G-F^* mice demonstrated significantly greater tumor number and area compared to wild type (Fig. 1, J and K). In contrast, AOM/DSS-treated female *App^NL-G-F^* mice showed significantly less tumors and area compared to their respective wild type mice (Fig. 1, L and M). Tumorigenesis was not observed in male and female vehicle-treated groups. AOM/DSS treatment did not induce colorectal-associated symptoms in Female *App^NL-G-F^* mice. These data indicate the AOM/DSS administration produced various sex- and genotype-dependent tumor numbers and areas.

### Histologic severity of tumorigenesis and inflammation induced by AOM/DSS treatment were sex- and APP-associated

We hypothesized that differential inflammatory responses might contribute to the sex- and genotype-dependent phenotypes. Exposure to AOM/DSS caused moderate to severe inflammation in both wild type and *App^NL-G-F^* male mice. (Fig. 2, A, C and F). The *App^NL-G-F^* treated with AOM/DSS exhibited significantly increased mucosal hyperplasia, neoplastic transformation, and higher overall histology scores compared to the wild type mice. Most of the *App^NL-G-F^* females did not show robust inflammation following AOM/DSS administration compared to the wild type groups. In contrast, wild type females showed drastically greater inflammation, mucosal hyperplasia, neoplastic transformation, and total histology scores compared to the *App^NL-G-F^*female AOM/DSS treated group (Fig. 2, B, G to J). These observations indicate varying susceptibilities to CAC based on genotype and sex, with *App^NL-G-F^* males exhibiting more severe colonic pathology compared to wild type mice, and *App^NL-G-F^* females showing less tumorigenesis compared to wild type groups. Notably, *App^NL-G-F^*males developed adenocarcinoma, while females showed histology from normal to mild in their colons.

**Fig. 2.**
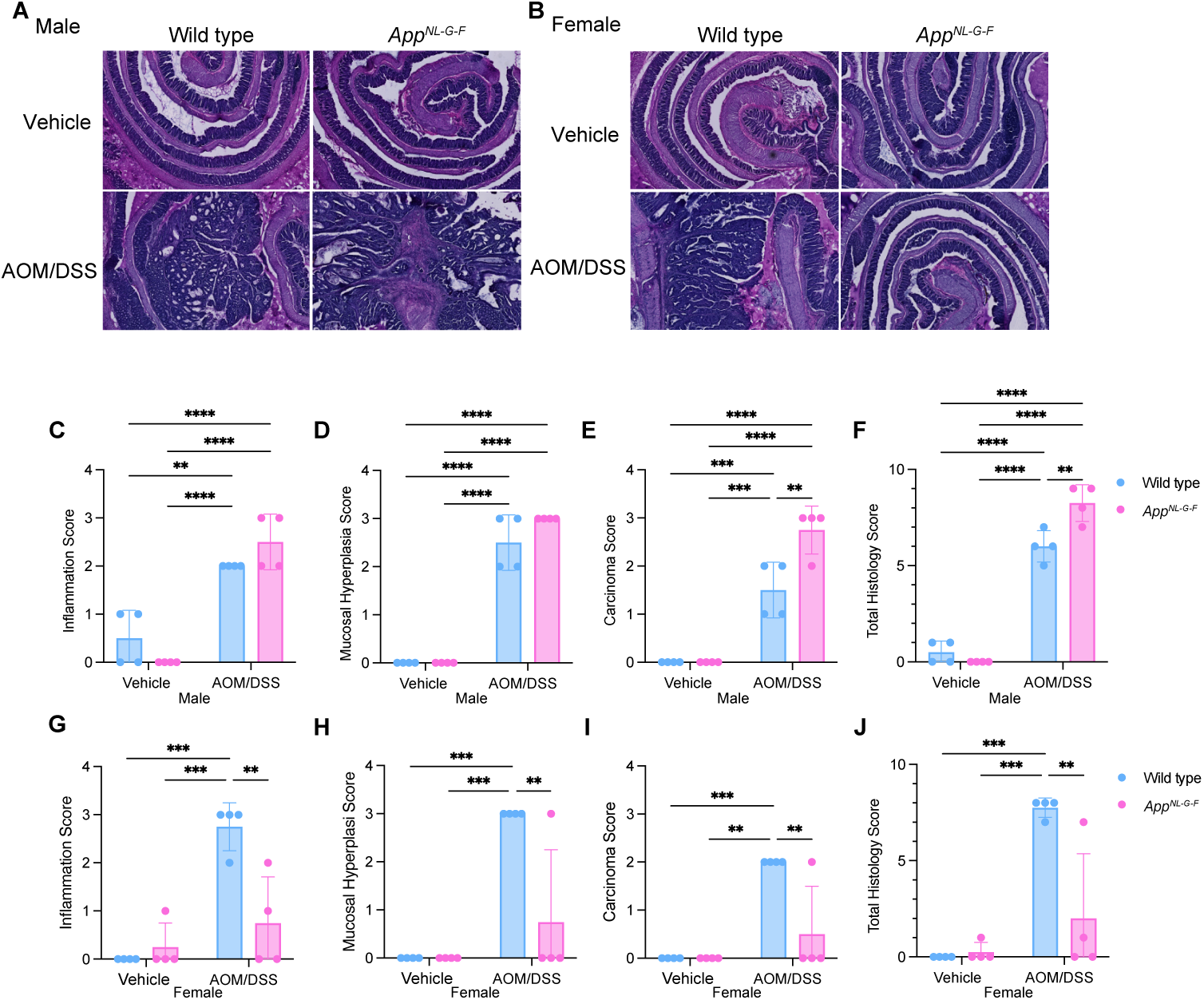
APP mutation reduced colon inflammation in AOM/DSS treated female mice. **A**,**B**, H&E staining of Swiss-rolls of normal colon and AOM/DSS-induced CAC are shown with 4X magnification. Histological lesions of wild type and *App^NL-G-F^* mouse colons are shown by scoring inflammation (**C**,**G**), mucosal hyperplasia (**D**,**H**), neoplastic transformation (**E**,**I**), and total histology (**F**,**J**), *p<0.05, ** p<0.01, ***p<0.001, and ****p<0.0001 (mean ± SEM, n=4 animals)

### AOM/DSS treated APP mutant mice exhibited sex dependent transcriptome changes in the colon

To understand why AOM/DSS treatment resulted in differential phenotypic outcomes in an APP mutant background at the molecular level, we conducted RNA-seq using colon cancer tissues, adjacent normal colon tissues, or normal colon tissues when tumors were not formed. The PCA plot clearly showed that gene expression profiles in tumors differed significantly from those in non-cancerous colon tissues, regardless of sex (Fig. 3A). Differential gene expression analysis using DESeq2 revealed significant impacts of *App^NL-G-F^* during AOM/DSS induced colon carcinogenesis (Fig. 3, B and C, Fig. S3A). For instance, in male mice, colon tumors from *App^NL-G-F^* mice (KMT) exhibited 1,672 up-regulated genes and 1,292 down-regulated genes (at FDR < 0.05) compared to colon tumors from wild type mice (WMT). Similarly, in female normal colon tissues, 1,741 up-regulated genes and 1,287 down-regulated genes were observed at FDR < 0.05 when comparing gene expression data between *App^NL-G-F^* (KFN) and wild type (WFN) mice. Even more substantial transcriptome differences were observed between normal colons from female *App^NL-G-F^*mice and colon tumors developed in female wild type mice. To explore whether APP mutations induced similar gene expression changes between male and female mice, differentially expressed genes (DEGs) were compared between datasets derived from males and females (Fig. 3D). While commonly up- or down-regulated genes were identified (920 genes and 541 genes, respectively), sex-dependent differences were evident from the DEG overlap analysis (Fig. 3D). To identify molecular pathways enriched in these DEGs, KEGG gene set enrichment analysis was performed using the iDEP2.0 online tool (Fig. 3E, Fig. S3, B and C). In both male and female comparisons between APP mutant and wild type strains, pathways related to steroid hormone biosynthesis and metabolism, such as nitrogen and retinol, were enriched among the DEGs. Cell cycle, TNF, and IL-17 signaling pathways were specifically enriched in female colons with APP mutations. Interestingly, although the *App^NL-G-F^* female mice did not develop colon tumors, genes associated with cancer development and mesenchymal-to-epithelial transition (EMT) were significantly altered when AOM/DSS-treated colon tissues in *App^NL-G-F^*female mice were compared with normal colon tissues from APP wild type mice (Fig. 3F). These results highlight the sex-dependent transcriptome differences in APP mutant colons during AOM/DSS-induced colitis.

**Fig. 3.**
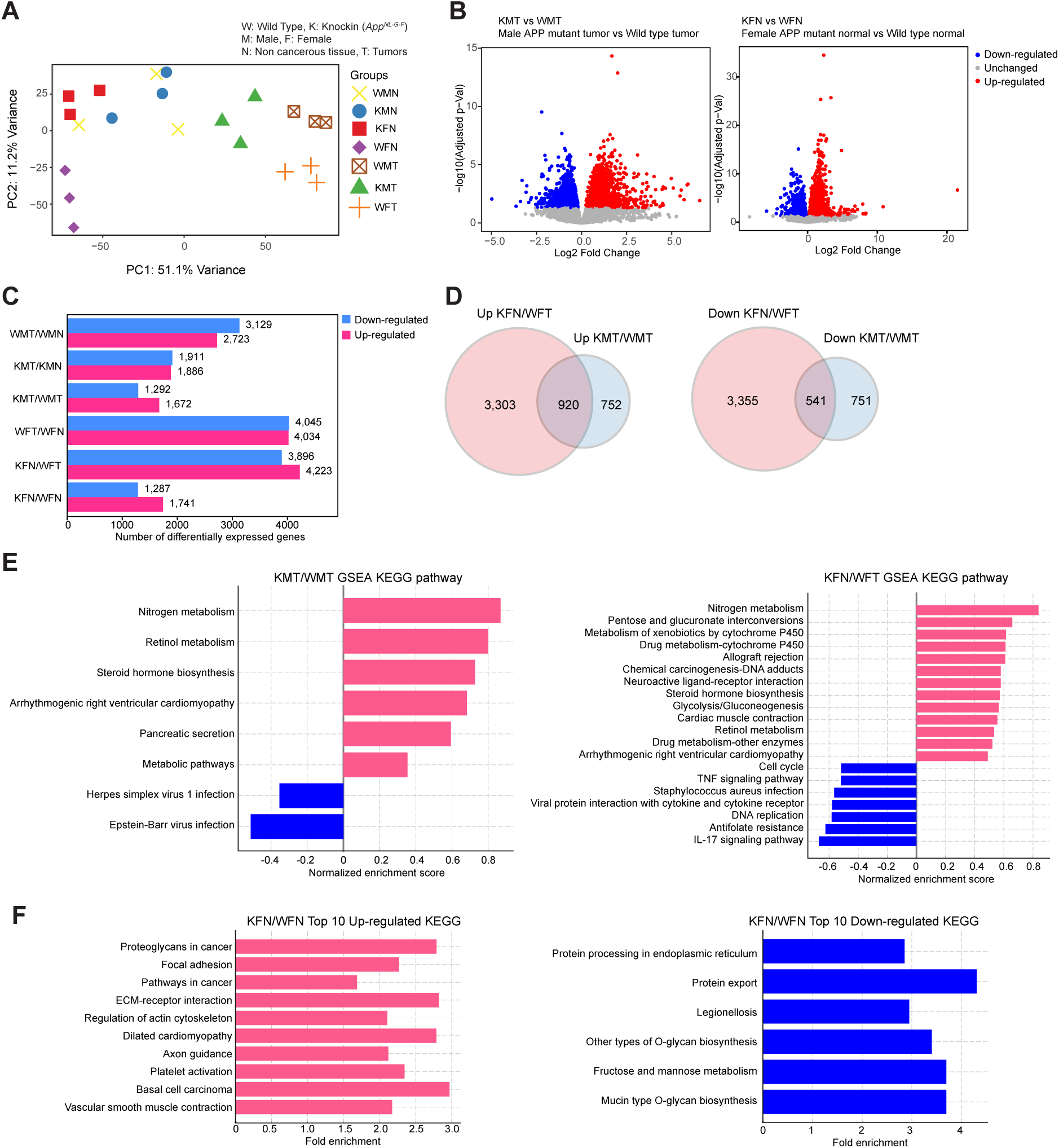
APP mutation led to sex dependent transcriptome alterations in the colon carcinogenesis model. **A,** Principal component analysis (PCA) of gene expression in colon cancer or normal colon tissues from AOM/DSS treated wild type and *App^NL-G7^*^09^*^F^* mice across different gender groups. The PCA plot depicts variance in gene expression profiles. Except for KFN (AOM/DSS treated female *App^NL-G-F^* normal tissue), in which tumors did not form, colon tumors and normal colon tissues were collected from the same mice in adjacent areas. RNA-seq data were obtained from three biological replicates. **B,** Volcano plots showing differential gene expression between male *App^NL-G-F^* tumors vs. wild type tumors (left panel) and between female *App^NL-G-F^* normal vs. wild type normal tissues (right panel). In both plots, red dots represent significantly up-regulated genes, blue dots indicate significantly down-regulated genes, and gray dots show genes with no significant change. Gene expression data from wild type tumors or normal tissues were used as a reference. **C,** Bar graphs of differentially expressed genes (DEGs) from various comparisons. The bars are colored to indicate up-regulated (red) and down-regulated (blue) genes, accompanied by the respective numbers of DEGs. **D,** Venn diagrams representing the overlap of up- or down-regulated genes in female *App^NL-G-F^* non-cancerous colons and male *App^NL-G-F^*colon tumors. Gene expression profiles of female wild type colon tumors and male wild type colon tumors were used as references for each comparison. **E,** Gene Set Enrichment Analysis (GSEA). Pathways significantly enriched (adjusted p-values < 0.05) in the comparison between male *App^NL-G-F^* tumors and wild type tumors (left panel), and female *App^NL-G-F^* normal tissues versus wild type normal tissues (right panel), are displayed with normalized enrichment scores. Pink bars represent positively enriched pathways, while blue bars denote negatively enriched pathways. **F,** Bar Charts of Top 10 KEGG Pathways. Top 10 up-regulated pathways in female *App^NL-G-F^* non-cancerous vs. wild type non-cancerous colon tissues are presented in the left panel, and down-regulated pathways are shown in the right panel, each with their respective fold enrichment scores.

### Differential expression of markers in M1 and M2 macrophages across various Genotypes

RNA-seq results suggested distinct stimulation of immune cell signaling pathways between males and females. Immune cells including macrophages are known to be recruited to tumor development sites^21^. Therefore, we examined whether AOS/DSS treatment resulted in differential immune responses. Consistent with this hypothesis, gene expression data revealed elevated levels of M1 marker gene expression in tumors from both male and female mice compared to noncancerous tissues (Fig. 4A), while M2 markers tended to be elevated in *App^NL-G-F^* female colons (Fig. 4B).

**Fig. 4.**
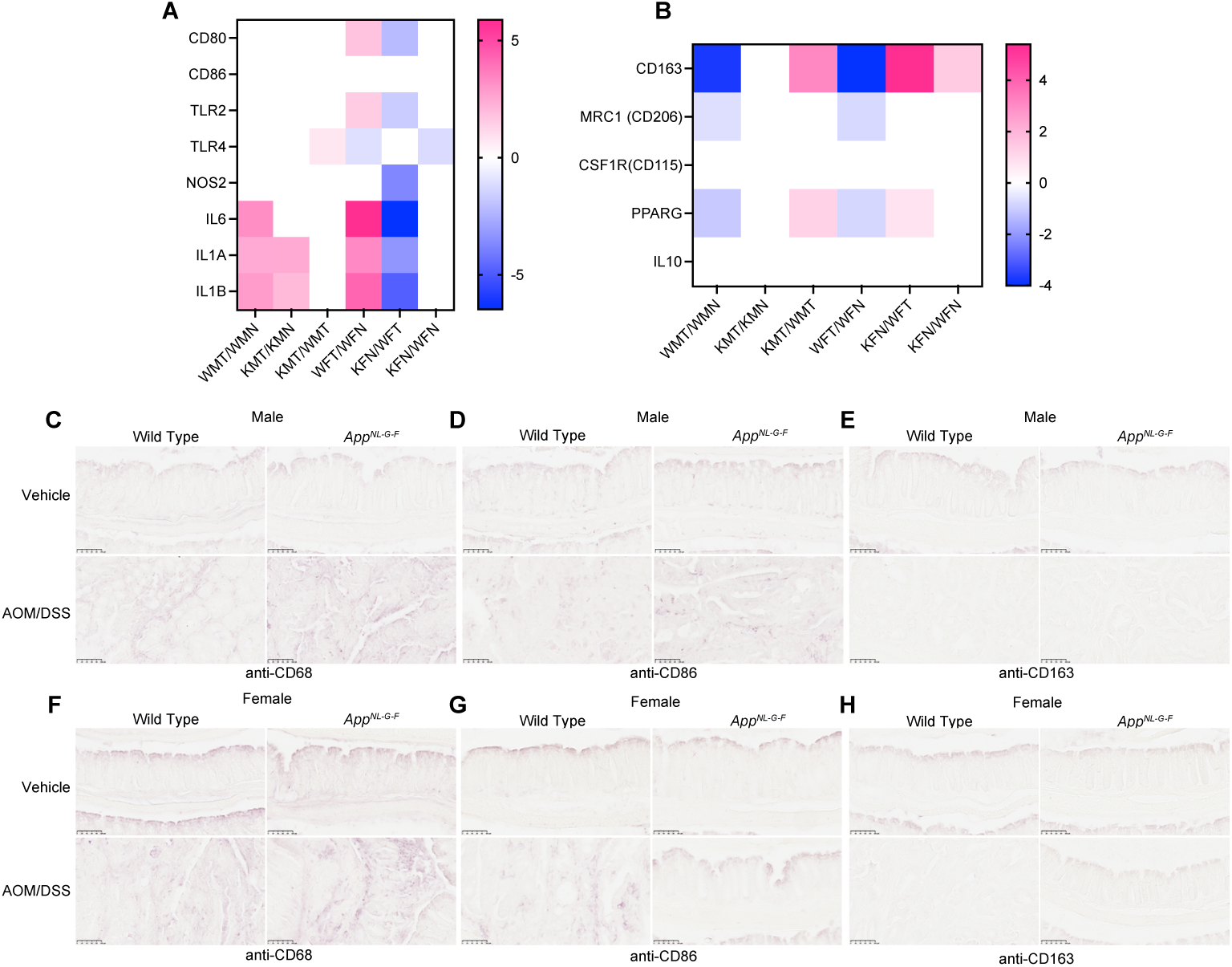
Tumor microenvironment was distinct between male and female *App^NL-G-F^* mice. **A**, Heatmap showing the expression of M1 macrophage markers (CD80, CD86, TLR2, TLR4, NOS2, IL6, IL1A, IL1B) across different experimental conditions. Pink indicates higher expression, blue indicates lower expression, with a scale ranging from -5 to 5. Columns represent various experimental groups, highlighting variations in gene expression. **B**, Heatmap showing the expression levels of M2 macrophage markers (CD163, MRC1 (CD206), CSF1R (CD115), PPARG, IL10) across different experimental conditions. Colors range from blue (lower expression) to pink (higher expression), with a scale from -4 to 4. The columns represent different experimental groups, illustrating variations in gene expression related to M2 macrophage phenotypes. **C**,**F** Immunohistochemical staining [n=8 replicates (4 biological replicates x 2 technical replicates, total 8 replicates)] for CD68 in colon tissues from male (**c**) and female (**f**) wild type and *App^NL-G-F^* mice treated with vehicle or AOM/DSS. The images from AOM/DSS are from cancerous tissue except for female *App^NL-G-F^* mice. **D**,**G**, Immunohistochemical staining for CD86 in colon tissues from male (**D**) and female (**G**) wild type and *App^NL-G-F^* mice treated with vehicle or AOM/DSS. **E**,**H**, Immunohistochemical staining for CD206 in colon tissues from male (**E**) and female (**H**) wild type and *App^NL-G-F^* mice treated with vehicle or AOM/DSS.

Immunohistochemical analyses also supported these findings, showing enhanced M1 macrophage staining in tumors from both genders. In contrast, CD163, a marker of M2 macrophages, exhibited similar levels regardless of APP status or AOS/DSS treatment (Fig. 4C). These results suggest that immune responses are uniquely modulated between male and female mice carrying APP mutations, including differential recruitment of macrophage subtypes. In general, the M1-type macrophages are considered pro-inflammatory and inhibitory for tumor growth^22^. The presence of M1 macrophage in our model suggests a distinct immune microenvironment. Consequently, *App^NL-G-F^* female mice may develop a unique protective immune microenvironment, shielding them from tumor development.

### Increased expression of neuronal markers in colons of female *App^NL-G-F^*mice

Many studies have demonstrated a functional link between neural responses and cancer progression^23,24^. Our RNA-seq analysis also indicated an upregulation of neuroactive ligand-receptor interactions in female colons with APP mutations (Fig. 3E). Specifically, expression levels of the neural marker PGP9.5 and the neural signaling enzyme Choline Acetyltransferase (ChAT) were elevated in the colons of *App^NL-G-F^* female mice compared to tumors from wild type female mice (Fig. 5A). In line with these findings, neurotransmitter receptors such as adrenergic, nicotinic, and muscarinic receptors were also up-regulated in the same experimental condition (Fig. 5, B and C). Immunohistochemical results further confirmed enhanced staining for PGP9.5, ChAT, and Tyrosine Hydroxylase (TH), particularly in *App^NL-G-F^* females treated with AOS/DSS (Fig. 5D). APP expression was confirmed in colon epithelial cells and entric neurons (Fig. S5). These results suggest potential roles for neural regulation in colon tumorigenesis, which may contribute to an antitumor function in the colon environment of female APP mutant mice.

**Fig. 5.**
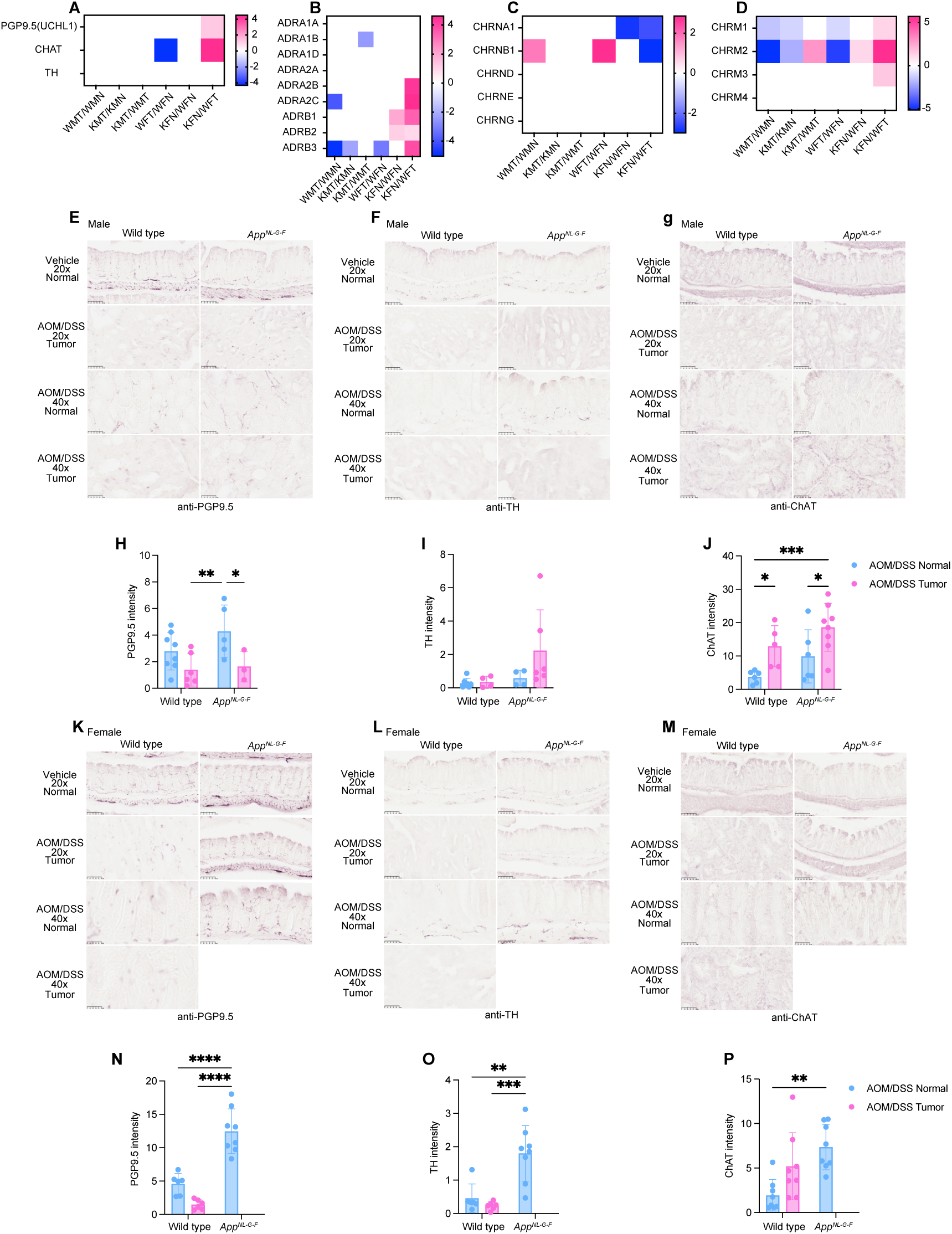
Neuronal markers were up-regulated in female *App^NL-G-F^* mice colons. **A - D,** Heatmap showing the expression levels of neuronal markers PGP9.5 (UCHL1), ChAT, TH (**A**), adrenergic receptors ADRA1A, ADRA1B, ADRA1D, ADRA2A, ADRA2B, ADRA2C, ADRB1, ADRB2, ADRB3 (**B**), neuronal nicotinic receptors CHRNA2, CHRNA3, CHRNA7, CHRNA9, CHRNA10, CHRNB2, CHRNB4 (**C**), and muscarinic receptors CHRM1, CHRM2, CHRM3, CHRM4 (**D**) across various experimental groups. The color scale from blue to pink indicates expression levels ranging from -4 (or -5) to 4 (or 5). Blue represents a decrease and pink an increase in marker expression. **E - G**, Histological analysis of colon tissue sections from male wild type and *App^NL-G-F^* mice, stained with anti-PGP9.5 (**E**), anti-Tyrosine Hydroxylase (TH, **F**), and anti-Choline acetyltransferase (ChAT, **G**) to visualize neuronal cells. The first row shows normal colon tissue from vehicle and treated mice at 20x magnification. The second and fourth rows show AOM/DSS-treated tissues at 20x and 40x magnifications, respectively, highlighting areas with tumor development. The third row provides a closer view (40x magnification) of normal colon tissue in the AOM/DSS treatment group. **H – J,** Quantification of the positive staining in male mice for PGP9.5, TH, and ChAT. *p<0.05, ** p<0.01, ***p<0.001, and ****p<0.0001 [(mean ± SEM, n=8 replicates(4 biological replicates x 2 technical replicates, total 8 replicates)]**K** – **M,** Histological analysis of colon tissue sections from female wild type and *App^NL-G-F^*mice, stained with anti-PGP9.5 (**K**), anti-TH (**L**), and anti-ChAT (**M**) to visualize neuronal cells. In the fourth row, no tumor images from AOM/DSS treated group as no tumor formation is observed. **N** – **O,** Quantification of the positive staining in female mice for PGP9.5, TH, and ChAT.

### CAC induced by AOM/DSS treatment affects brain Aβ Accumulation in both *App^NL-G-F^* Males and Females

Collectively, our results demonstrate that mutant APP expression alters the colonic tumorigenic response following AOM/DSS treatment. We have previously identified a crosstalk between the colon and brain, showing that DSS-induced colitis-like conditions exacerbate brain Aβ pathology^25,26^. Although we observed no detectable Aβ immunoreactivity in the colons (Fig. S5B) we investigated whether AOM/DSS-induced tumorigenesis similarly affects brain pathology. The impact of CAC induced by AOM/DSS on Aβ accumulation in the brain was assessed by ELISA. Results showed that soluble Aβ 1-40 levels decreased in AOM/DSS treated male *App^NL-G-F^* mice compared to the vehicle group (Fig. 6A), although the total Aβ immunoreactivity in male *App^NL-G-F^* brains were increased upon AOM/DSS treatment (Fig. S6A). Interestingly, in AOM/DSS treated female *App^NL-G-F^* mice, levels of insoluble Aβ 1-40, soluble Aβ 1-42, and insoluble Aβ 1-42 were also decreased compared to vehicle (Fig. 6B).

**Fig. 6.**
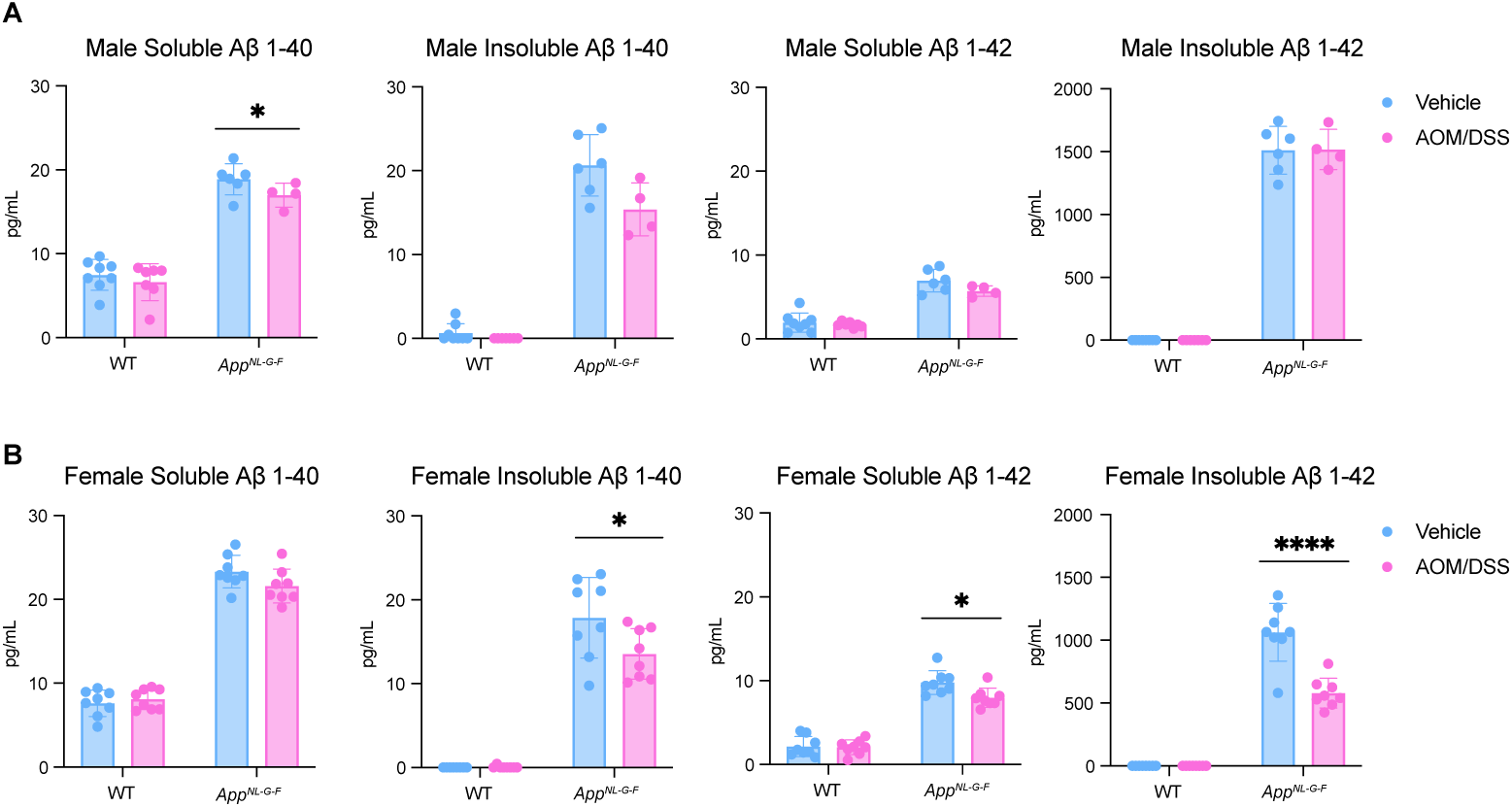
Aβ 1-42 levels were reduced in AOM/DSS treated female *App^NL-G-F^* brains. **A**,**B,** Measurement of soluble and insoluble Aβ 1-40 and Aβ 1-42 in male (**A**) and female (**B**) wild type and *App^NL-G-F^* brains (partial cortex) from mice treated with either vehicle or AOM/DSS. *p<0.05, ** p<0.01, ***p<0.001, and ****p<0.0001 (mean ± SEM, n=4-8 replicates)

Microglia and astrocytes, play critical roles in the hypothesized pathophysiology of AD^27^. To observe glia changes, differences in immunoreactivity were quantified. Iba1 immunostaining was increased in AOM/DSS treated male wild type mice, but was decreased in AOM/DSS treated *App^NL-G-F^* males when compared to their respective vehicle groups (Fig. S6, A to C). However, both wild type and *App^NL-G-F^* females exhibited increased GFAP immunoreactivity following treatment. In summary, these findings suggest that the inflammatory and/or carcinogenic processes triggered by AOM/DSS may interact with amyloidogenic and gliosis pathways in a sex dependent manner.

## Discussion

The number of Alzheimer’s disease (AD) cases in the U.S. is projected to rise significantly, potentially affecting 14 million people by 2050^28^, which nearly doubles the current figures. Similarly, global cancer incidence and mortality are expected to see a substantial increase, with cancer-related deaths projected to rise by 89.7% by 2050 compared to 2022 levels^29^. Epidemiological studies have highlighted that AD patients generally have a lower risk of developing cancer, and conversely, cancer patients exhibit reduced risks of AD^30^. However, the biological mechanisms underlying this inverse relationship remain unclear. In this study, we induced CAC in the *App^NL-G-F^* amyloidosis AD mouse model to investigate the effects of sex and genotype on cancer development and to examine how the AD condition influences CAC pathological outcomes.

Surprisingly, our data showed that tumor formation was inhibited in female *App^NL-G-F^* mice but stimulated in *App^NL-G-F^*male mice (Fig. 1). This suggests a sex-specific influence of APP mutations on tumorigenesis. We confirmed APP expression in both normal colons and human colon cancers across various grades of differentiation (Fig. 1, A and B). Clinical data further suggest differential roles of APP in the outcomes of human colorectal cancer patients. While high APP expression is associated with worse overall survival in both male and female patients, relapse-free rates are significantly correlated with higher APP expression in males, whereas in females, the opposite trend is observed (Fig. 1, C and D and Fig. S1, A and B). APP and its AD-causing mutations may play an active role in tumor progression and patient prognosis, particularly in males, who exhibit poorer survival with higher APP levels.

The sex-specific responses to AOM/DSS treatment, particularly the increased tumor burden in male *App^NL-G-F^*mice compared to females, might be influenced by differences in inflammatory responses and APP-mediated pathways. The exacerbated inflammatory and tumorigenic responses in male mice (Fig. 2, A, C to F) compared to the milder responses in female mice could be a result of different interactions between APP mutations and sex hormones or other genetic factors that modulate inflammation and cellular proliferation. The differences in inflammation severity and tumor development between sexes (Fig. 2, B, G to J) further suggests a genotype-dependent susceptibility to CAC. The marked histological differences indicate that male *App^NL-G-F^* mice are more prone to severe colitis and subsequent neoplastic transformation, possibly due to a more aggressive inflammatory milieu driven by APP mutations.

The distinct transcriptomic profiles observed in male and female mice (Fig. 3), particularly in APP mutant mice, highlight the influence of APP mutations on gene expression changes during colon carcinogenesis. The identification of sex-dependent pathways, such as those involved in steroid hormone biosynthesis and inflammatory signaling, suggests that different biological mechanisms are preferentially activated in males and females in response to APP mutations and AOM/DSS treatment. The differential expression of immune markers, specifically M1 and M2 macrophage markers in APP mutant mice (Fig. 4), suggests a unique immune environment that may contribute to the sex-specific tumor outcomes observed. Since M1 macrophages are generally considered anti-tumoral^31^, this finding suggests a complex interaction between AD pathology and the colon microenvironment. Female mice, on the other hand, displayed a distinct immune profile or tumor microenvironment. The increased expression of neuronal markers in the colons of female APP mutant mice (Fig. 5) suggests an intriguing link between neural regulation and tumor resistance. This could imply a neuroimmune interaction where neuronal signaling influences immune responses and cancer prohibition. We observed upregulation of the β3-adrenergic receptor (ADRB3) and the α2 subunit of the nicotinic acetylcholine receptor (CHRNA2) in female *App^NL-G-F^* mice treated with AOM/DSS, in contrast to similar wild type female mice, with no corresponding changes in males. β3-adrenergic receptors are known to be involved in modulating inflammatory responses and fat metabolism, both key factors in cancer development^32^. These receptors may inhibit cellular pathways that promote proliferation and survival, providing a mechanistic basis for their involvement in disease modulation. Moreover, this receptor upregulation could represent a compensatory mechanism to altered catecholamine dynamics, which are known to be disrupted in AD, affecting neurotransmitter levels and receptor expression.

Similarly, CHRNA2 upregulation could impact cellular processes such as proliferation, apoptosis, and angiogenesis, influencing both tumor development and the neurological alterations associated with AD^33,34^. The specific enhancement of CHRNA2 expression in females suggests that estrogen or other sex-specific factors may enhance the expression or functional activity of these receptors in response to the physiological stresses imposed by AOM/DSS treatment. Additionally, the observed decrease in Aβ40 and Aβ42 levels in the brains of AOM/DSS-treated *App^NL-G-F^* female mice compared to vehicle treated *App^NL-G-F^* female mice, along with the upregulation of ADRB3 and CHRNA2, provides a complex picture of how systemic inflammation and local neurotransmitter signaling may interact with Alzheimer’s disease-related pathology. This decrease in Aβ peptides, potentially indicative of enhanced clearance or reduced production, could be an adaptive response to mitigate the dual challenges of neurodegeneration and systemic inflammation, further influenced by the modulation of adrenergic and nicotinic receptor pathways. Both ADRB3 and CHRNA2 are integral to networks involving not only their primary ligands but also crosstalk with other signaling pathways, including those mediated by other neurotransmitters and catecholamines^33^. This complex interaction network likely forms part of a broader adaptive response to the dual challenges of neurodegeneration and potential carcinogenesis presented by AD pathology and AOM/DSS exposure.

Interestingly, despite the absence of cancer formation, several cancer pathways were upregulated in female APP mutant mice treated with AOM/DSS (Fig.1I and Fig. 3F). This could indicate a potential mechanism for preventing tumorigenesis, suggesting that the APP mutations may confer a unique protective effect in these mice. Understanding these mechanisms could provide insights into new preventative strategies for colorectal cancer, particularly in populations at high risk due to genetic factors. The activation of cancer-related pathways without subsequent tumor formation raises several intriguing possibilities. It may suggest that despite the activation of oncogenic signaling, additional checks and balances within the cellular environment of these APP mutant females might inhibit the final transformation steps necessary for cancer development. This could be due to enhanced DNA repair mechanisms, apoptosis, or cellular senescence processes that are more active or efficient in these female mice. Alternatively, the upregulation of these pathways might reflect a state of heightened cellular defense against tumorigenesis, potentially mediated by immune surveillance mechanisms that are more robust in these particular genetic backgrounds.

In conclusion, our findings suggest that APP plays a multifaceted role in CAC, with significant implications for understanding the biological links between CAC and AD. These results emphasize the importance of considering sex and genetic background in the study and treatment of CAC and potentially AD, paving the way for more personalized therapeutic approaches.

## Methods

### Animals

Male and female wild-type C57BL/6 mice and APP knock-in transgenic mice (*App^NL-G-F^*, KI:RBRC06344) were used in this study. Wild-type mice were originally purchased from The Jackson Laboratory (Bar Harbor, Maine), while the *App^NL-G-F^* mice were generously provided by Dr. Takashi Saito and Dr. Takaomi C. Saido from the RIKEN BioResource Center, Japan^19^. The *App^NL-G-F^*model features a humanized Aβ sequence carrying the Swedish (NL), Arctic (G), and Beyreuther/Iberian (F) mutations. These mutations respectively enhance Aβ production, promote aggregation by facilitating oligomerization and reducing proteolytic degradation, and increase the Aβ42/40 ratio^19^.

All mice were maintained at the University of North Dakota Center for Biomedical Research. Mice were housed in 12 h light/dark cycle with food and water provided *ad libitum*. For the experiment, 6-11 male and female mice per treatment group, aged 5-8 months, were used. The mice were randomly assigned to either a vehicle group (saline via intraperitoneal [i.p.] injection and drinking water) or a drug treatment group (azoxymethane [AOM] via i.p. injection and dextran sodium sulfate [DSS] in drinking water). At the end of the study, mice were euthanized, and brain, spleen, and colon were collected for histological and biochemical analyses. All animal procedures were reviewed and approved by the University of North Dakota Institutional Animal Care and Use Committee (UND IACUC) and adhered to the *Guide for the Care and Use of Laboratory Animals* (8th Edition, National Research Council of the National Academies).

### Induction of colitis-associated colorectal cancer (CAC) via AOM/DSS Treatment

To mimic human colorectal cancer pathology, mice received a 10 mg/kg i.p. injection of AOM (Sigma-Aldrich, St. Louis, MO, USA) on day 0 (week 0) and were then exposed to 1.5% w/v DSS (MW = 36–50 kDa, MP Biomedicals, LLC, Santa Ana, CA, USA) in autoclaved drinking water starting in week 1 for a duration of 7 days. Control (vehicle-treated) groups received only saline via i.p. injections on day 0 and had continuous access to autoclaved drinking water throughout the entire study period^35–37^. During the DSS exposure, DSS consumption was recorded per mouse per day to ensure consistent intake. Throughout the 17-week experimental timeline, body weights were recorded weekly to monitor treatment-related changes, except during week 1 when animals were housed in isolated chambers due to the carcinogenic nature of AOM. Percent body weight change was calculated weekly for each animal by dividing the current weight by its initial weight at week 0 and expressing the result as a percentage. At the end of the 17-week period, animals were euthanized. The colon length from the cecum to the rectum and the weight of the colon and spleen were measured for each animal. The number of visible tumors was counted along the entire colon, from proximal to distal segments. Tumor area (in mm²) was quantified for each mouse by calculating the average tumor size using Adobe Photoshop CS3 software (Adobe Systems, San Jose, CA) from high-resolution images captured at the time of tissue collection. For subsequent histological and biochemical analyses, tissues including the brain, spleen, and both normal and cancerous tissues of colon were collected and preserved accordingly.

### Histological Staining and Scoring of Colonic Swiss-rolls

The entire colon, from proximal to distal segments, was collected and processed into Swiss-rolls^38^. Tissues were fixed in 4% paraformaldehyde (PFA) for five days and cryoprotected by two changes in 30% sucrose before being embedded in 15% gelatin blocks. These blocks were then serially cryosectioned at a thickness of 10 μm. Hematoxylin and eosin (H&E)-stained sections were evaluated by a pathologist blinded to the experimental groups. CAC scores were assigned based on three histopathological parameters: inflammation, mucosal hyperplasia, and neoplastic transformation. Inflammation was scored on a 0–3 scale, 0= normal histology; 1= mild, focal, or scattered inflammation limited to the lamina propria; 2= moderate inflammation that was multifocal or locally extensive and extended into the submucosa; 3= severe, transmural inflammation. Mucosal hyperplasia was also scored on a 0–3 scale, 0= indicated normal crypt architecture; 1= mild hyperplasia with crypts 2–3 times thicker than normal while retaining normal epithelial morphology; 2= moderate hyperplasia with crypt elongation (2–3×), hyperchromatic epithelial nuclei, reduced goblet cell numbers, and scattered crypt branching (arborization); and 3= severe hyperplasia with crypts ≥ four times normal thickness, marked nuclear hyperchromasia, little to no goblet cells, a high mitotic index, and frequent arborization. Neoplastic transformation was scored as follows: 0= normal mucosa; 1= low-grade dysplasia characterized by mild nuclear atypia and preservation of glandular architecture; 2= high-grade dysplasia or adenomatous polyp with pronounced cellular atypia, crowding, and loss of polarity; and 3= adenocarcinoma showing invasive growth, glandular disorganization, and marked cytological abnormalities. Both individual parameter scores and their combined totals were plotted for each animal to provide a comprehensive evaluation of colonic pathology in the Swiss-roll preparations.

For CD86, CD163 and ChAT staining, Tris-EDTA pH 9.0 buffer was used at 95 °C for 10 min for antigen retrieval. For TH staining, antigen retrieval was performed using sodium citrate pH 6.0 at 95 °C for 10 min. The Swiss-rolls sections were blocked in IHC solution [0.5% bovine serum albumin (BSA, Equitech-Bio, Inc.), 0.1% Triton X-100 (Sigma-Aldrich), 5% normal goat serum (NGS, Equitech-Bio, Inc.), and 0.02% Na Azide] for 1 hour after blocking endogenous peroxidase activity using 3% H_2_O_2_ buffer for 5 mins. Subsequently, the sections were incubated with primary antibody: APP (1:500, ab32136, Abcam, Cambridge, MA, USA), TACE (1:1000, PA5-27395, Invitrogen, ThermoFisher Scientific, Waltham, MA, USA), PS1 (1:500, PA5-119872, Invitrogen, ThermoFisher Scientific, Waltham, MA, USA), BACE (1:1000, #5606, Cell Signaling Technology, Inc., Danvers, MA, USA), CD68 (1:1000, MCA1957, Bio-Rad Laboratories, Inc., CA, USA), CD86 (1:675, #19589, Cell Signaling Technology, Inc., Danvers, MA, USA), CD163 (1:500, ab182422, Abcam, Cambridge, MA, USA), PGP9.5 (1:1000, ab108986, Abcam, Cambridge, MA, USA), ChAT (1:2000, ab178850, Abcam, Cambridge, MA, USA), and TH (1:3000, ab137869, Abcam, Cambridge, MA, USA) at 4°C overnight. On the following day, tissues were incubated using biotinylated secondary antibody for 2 h. To visualize the antibody binding, a VECTASTAIN Avidin-Biotin Complex (ABC) kit was used followed by the Vector VIP Peroxidase (HRP) Substrate kit (SK-4600) (Vector Laboratories, Inc., Burlingame, CA, USA).

Frozen human colon tissue array slides containing both normal and tumor specimens were obtained from BioChain (Newark, CA, USA). H&E staining, and immunohistochemical analysis of APP expression was performed using anti-APP Y188 antibody (1:1000 dilution, Abcam, Cambridge, MA, USA). All stained slides were digitally scanned using a Hamamatsu NanoZoomer 2.0HT slide scanner.

### RNA-seq analysis (Colon tumor tissue)

Total RNAs were purified using Qiazol RNA Isolation Reagent (QIAGEN, Germantown, MD). Total RNA integrity was determined using Agilent Bioanalyzer or 4200 Tapestation. RNA samples were submitted for library preparation and sequencing to the Genome Technology Access Center at the McDonnell Genome Institute (GTAC@MGI), Washington University School of Medicine. Library preparation was performed with 500ng to 1ug of total RNA. Ribosomal RNA was removed by an RNase-H method using RiboErase kits (Kapa Biosystems). mRNA was then fragmented in reverse transcriptase buffer and heating to 94 degrees for 8 minutes. mRNA was reverse transcribed to yield cDNA using SuperScript III RT enzyme (Life Technologies, per manufacturer’s instructions) and random hexamers. A second strand reaction was performed to yield ds-cDNA. cDNA was blunt ended, had an A base added to the 3’ ends, and then had Illumina sequencing adapters ligated to the ends. Ligated fragments were then amplified for 12-15 cycles using primers incorporating unique dual index tags. Fragments were sequenced on an Illumina NovaSeq-6000 using paired end reads extending 150 bases. Basecalls and demultiplexing were performed with Illumina’s bcl2fastq software with a maximum of one mismatch in the indexing read. Filtered read pairs were mapped to the mm10 genome using HISAT2^39^(version 2.1.0). Gene-level read counts were obtained using Subread featureCounts (version 1.5.0-0-p1). Differential gene expression analysis, figure generation, and pathway enrichment analysis were performed using iDEP2.0^40,41^. Pathway analysis was conducted using Parametric Gene Set Enrichment Analysis (PGSEA)^42^ on KEGG gene sets, incorporating the fold change values of all genes. The top 10 upregulated and downregulated pathways were identified through KEGG pathway analysis, using DEGs as inputs.

### Brain Immunohistochemistry

The left brain was fixed in 4% PFA for five days. After fixation, tissues were cryoprotected through two changes in 30% sucrose and subsequently embedded in 15% gelatin blocks. The embedded brains were then serially sectioned at 40 μm thickness using a sliding microtome^43^. Antigen retrieval was required for Aβ and Iba1 immunostaining. For Aβ, the brain sections were incubated in 25% formic acid for 25 minutes at room temperature. For Iba1, Tris-EDTA pH 9.0 buffer was used at 95 °C for 10 min. Sections were blocked for at least 30 minutes in an IHC blocking solution containing 0.5% bovine serum albumin (BSA, Equitech-Bio, Inc.), 0.1% Triton X-100 (Sigma-Aldrich), 5% normal goat serum (NGS, Equitech-Bio, Inc.), and 0.02% sodium azide, and were then incubated overnight at 4°C with a primary antibody targeting Aβ (1:500, #8243, Cell Signaling Technology, Danvers, MA, USA), Iba1 (1:1000, rabbit, 019–19741; Wako Chemicals USA, Inc., Richmond, VA, USA) and GFAP (1:1000, rabbit, D1F4Q; Cell Signaling Technology, Inc., Danvers, MA, USA). The next day, sections were incubated with a biotinylated secondary antibodyfor 2 hours at room temperature. Signal detection was performed using the VECTASTAIN Avidin-Biotin Complex (ABC) kit, and signals were visualized using the Vector VIP Peroxidase (HRP) Substrate Kit (SK-4600, Vector Laboratories). Stained brain sections were imaged using a Hamamatsu NanoZoomer 2.0HT digital slide scanner. For quantification of Aβ-positive regions in the hippocampus, 2–3 representative hippocampal images from 4–10 animals per experimental group were analyzed using QuPath (version 0.4.4)^44^.

### ELISA for soluble and insoluble Aβ 1–40 and Aβ 1–42

Soluble and insoluble Aβ 1-40 and Aβ 1-42 were analyzed using an ELISA kit (DAB140 and DAB142) from R&D systems (Minneapolis, MN, USA) according to the manufacturer’s protocol. The partial cortex tissues, previously flash-frozen, were weighed and lysed in ice-cold RIPA buffer composed of 20 mM Tris-HCl (pH 7.4), 150 mM NaCl, 1 mM Na3VO4, 10 mM NaF, 1 mM EDTA, 1 mM EGTA, 0.2 mM phenylmethylsulfonyl fluoride, 1% Triton-X100, 0.1% SDS, and 0.5% deoxycholate, along with a cocktail of protease inhibitors including 1 mM AEBSF, 0.8 μM aprotinin, 21 μM leupeptin, 36 μM bestatin, 15 μM pepstatin A, and 14 μM E-64. Homogenization of the tissues was performed using Zirconium oxide beads in a Bullet Blender, followed by centrifugation to separate the soluble from the insoluble components. The supernatant was utilized for soluble Aβ 1-40 and Aβ 1-42 ELISA, while protein concentrations were determined using the BCA assay from Thermo Fisher Scientific. The pellet containing insoluble material was resuspended in 5 M guanidine HCl with 50 mM Tris-HCl (pH 8.0), re-homogenized, and centrifuged to obtain the supernatant for insoluble Aβ ELISA.

### Statistical Analysis

Data were analyzed using two-way ANOVA with multiple comparisons, followed by Uncorrected Fisher’s LSD tests, utilizing GraphPad Prism 10 software. Results are presented as mean ± SEM. Significance levels are denoted by P values, with P < 0.05 being considered significant. The levels of significance are marked as follows: *P < 0.05, **P < 0.01, ***P < 0.001, ****P < 0.0001.

## Acknowledgement

We gratefully thank the University of North Dakota School of Medicine and Health Sciences for providing research environment and student stipend support. We appreciate Dr. Motoki Takaku for constructive feedback on RNA-seq data analyses.

## Author contributions

Conceptualization: T.I.T., M.S., C.K.C.; Methodology: T.I.T., M.S., C.K.C.; Investigation: T.I.T, M.S., H.L.P., C.K.C.; Visualization: T.I.T.; Supervision: C.K.C., Writing—original draft: T.I.T., Writing—review & editing: T.I.T, M.S., H.L.P., C.K.C.

## Data and materials availability

RNA-seq raw and processed data have been deposited in Gene Expression Omnibus (GEO) under accession number GSE297490. Link for reviewers: https://www.ncbi.nlm.nih.gov/geo/query/acc.cgi?acc=GSE297490 Token: mbetqewgrzsrrux

