## Supplemental figures for "Alzheimer’s Disease-Related Mutations in APP Influence Colorectal Tumor Formation in a Sex-Dependent Manner"

Fig. S1.

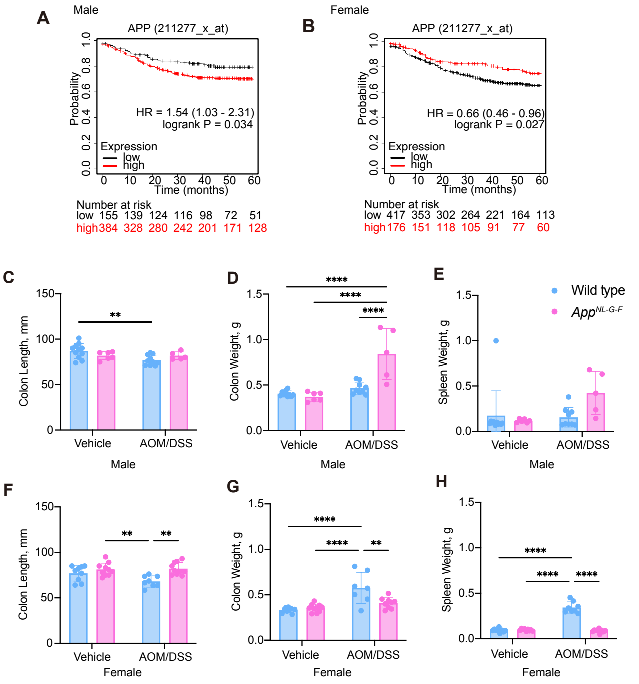


**Fig. S1. Comprehensive analysis of APP expression and physiological changes in response to AOM/DSS Treatment. A**,**B**, Kaplan-Meier curves for Relapse-Free Survival (RFS) in male (**A**) and female (**B**) subjects based on APP gene expression. The numbers below each plot indicate subjects at risk over 60 months, decreasing from initial counts. **C** – **H,** Graphs show colon length, colon weight, and spleen weight in male (**C** - **E**) and female (**F** - **H**) wild type and *App*$^{NL\text{-}G\text{-}F}$ mice treated with either vehicle or AOM/DSS.

Fig. S3-1.

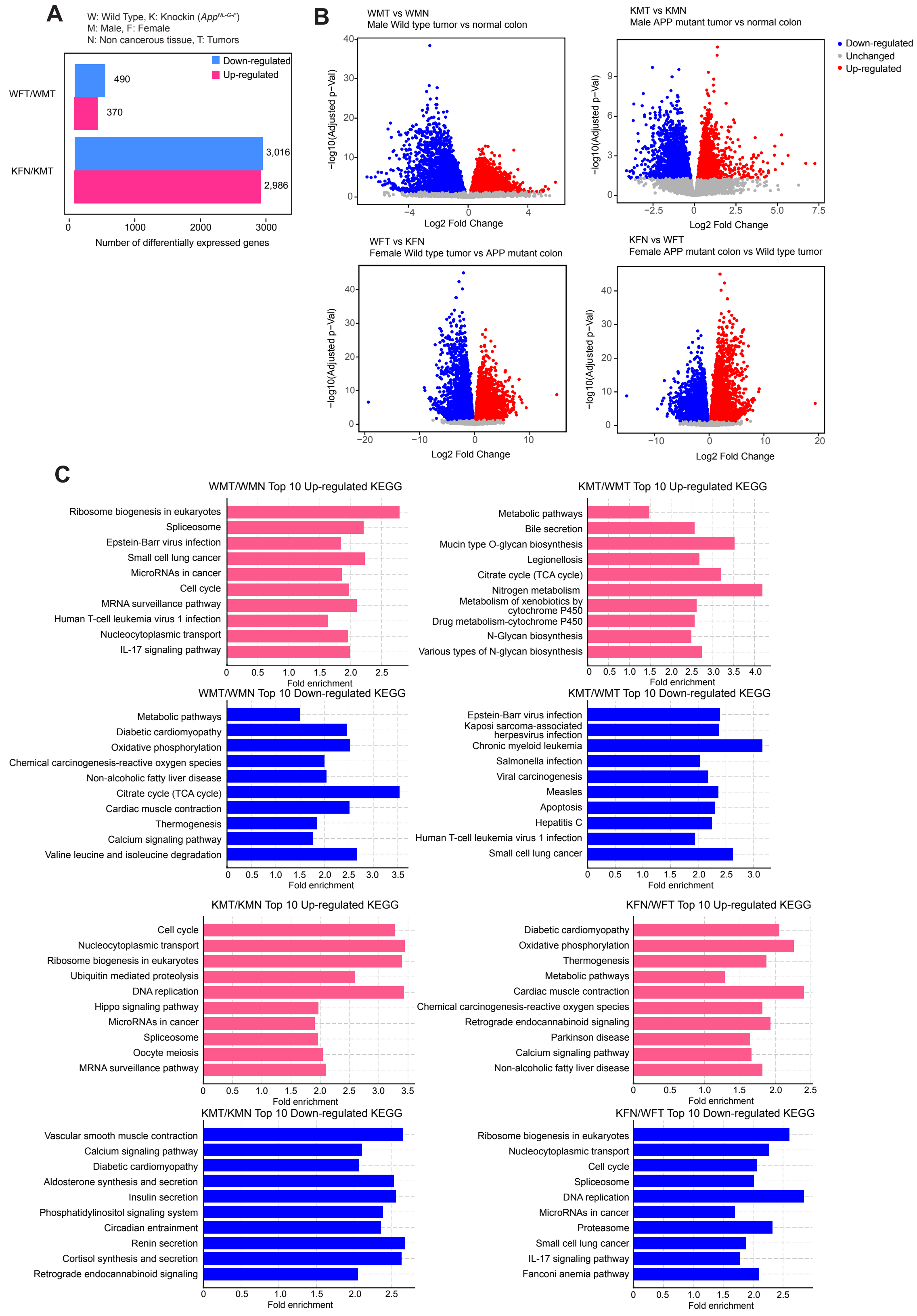

A
W: Wild Type, K: Knockin (App^NL-G-F)
M: Male, F: Female
N: Non cancerous tissue, T: Tumors
Down-regulated
Up-regulated
WFT/WMT
KFN/KMT
490
370
3,016
2,986
Number of differentially expressed genes
B
WMT vs WMN
Male Wild type tumor vs normal colon
KMT vs KMN
Male APP mutant tumor vs normal colon
WFT vs KFN
Female Wild type tumor vs APP mutant colon
KFN vs WFT
Female APP mutant colon vs Wild type tumor
Down-regulated
Unchanged
Up-regulated
−log10(Adjusted p−Val)
Log2 Fold Change
C
WMT/WMN Top 10 Up-regulated KEGG
Ribosome biogenesis in eukaryotes
Spliceosome
Epstein-Barr virus infection
Small cell lung cancer
MicroRNAs in cancer
Cell cycle
MRNA surveillance pathway
Human T-cell leukemia virus 1 infection
Nucleocytoplasmic transport
IL-17 signaling pathway
Fold enrichment
KMT/WMT Top 10 Up-regulated KEGG
Metabolic pathways
Bile secretion
Mucin type O-glycan biosynthesis
Legionellosis
Citrate cycle (TCA cycle)
Nitrogen metabolism
Metabolism of xenobiotics by cytochrome P450
Drug metabolism-cytochrome P450
N-Glycan biosynthesis
Various types of N-glycan biosynthesis
WMT/WMN Top 10 Down-regulated KEGG
Metabolic pathways
Diabetic cardiomyopathy
Oxidative phosphorylation
Chemical carcinogenesis-reactive oxygen species
Non-alcoholic fatty liver disease
Citrate cycle (TCA cycle)
Cardiac muscle contraction
Thermogenesis
Calcium signaling pathway
Valine leucine and isoleucine degradation
KMT/WMT Top 10 Down-regulated KEGG
Epstein-Barr virus infection
Kaposi sarcoma-associated herpesvirus infection
Chronic myeloid leukemia
Salmonella infection
Viral carcinogenesis
Measles
Apoptosis
Hepatitis C
Human T-cell leukemia virus 1 infection
Small cell lung cancer
KMT/KMN Top 10 Up-regulated KEGG
Cell cycle
Nucleocytoplasmic transport
Ribosome biogenesis in eukaryotes
Ubiquitin mediated proteolysis
DNA replication
Hippo signaling pathway
MicroRNAs in cancer
Spliceosome
Oocyte meiosis
MRNA surveillance pathway
KFN/WFT Top 10 Up-regulated KEGG
Diabetic cardiomyopathy
Oxidative phosphorylation
Thermogenesis
Metabolic pathways
Cardiac muscle contraction
Chemical carcinogenesis-reactive oxygen species
Retrograde endocannabinoid signaling
Parkinson disease
Calcium signaling pathway
Non-alcoholic fatty liver disease
KMT/KMN Top 10 Down-regulated KEGG
Vascular smooth muscle contraction
Calcium signaling pathway
Diabetic cardiomyopathy
Aldosterone synthesis and secretion
Insulin secretion
Phosphatidylinositol signaling system
Circadian entrainment
Renin secretion
Cortisol synthesis and secretion
Retrograde endocannabinoid signaling
KFN/WFT Top 10 Down-regulated KEGG
Ribosome biogenesis in eukaryotes
Nucleocytoplasmic transport
Cell cycle
Spliceosome
DNA replication
MicroRNAs in cancer
Proteasome
Small cell lung cancer
IL-17 signaling pathway
Fanconi anemia pathway

**Fig. S3-1. Comprehensive comparisons of gene expression data from *App*$^{NL\text{-}G\text{-}F}$ normal colon tissues and tumors. A,** Bar graphs showing sex dependent differences in colon gene expression data. Bars are colored to indicate up-regulated (pink) and down-regulated (blue) genes, accompanied by the respective numbers of DEGs. **B,** Volcano plots showing DEGs in various comparisons. Red or blue dots represent significantly up- or down-regulated genes, while gray dots show genes with no significant changes. Top left (WMT vs WMN): Male wild type tumor vs. normal colon. Top right (KMT vs KMN): Male *App*$^{NL\text{-}G\text{-}F}$ tumor vs. normal colon. Bottom left (WFT vs KFN): Female wild type tumor vs. *App*$^{NL\text{-}G\text{-}F}$ normal colon. Bottom right (KFN vs WFT): Female *App*$^{NL\text{-}G\text{-}F}$ normal colon vs. wild type tumor. **C,** Bar charts showing the top 10 KEGG pathways. The top 10 KEGG pathways enriched in DEGs identified in part (b) are displayed. Blue bars indicate down-regulated pathways, while pink bars represent up-regulated pathways, each ranked by their fold enrichment scores. RNA-seq data were obtained from three biological replicates.

Fig. S3-2.

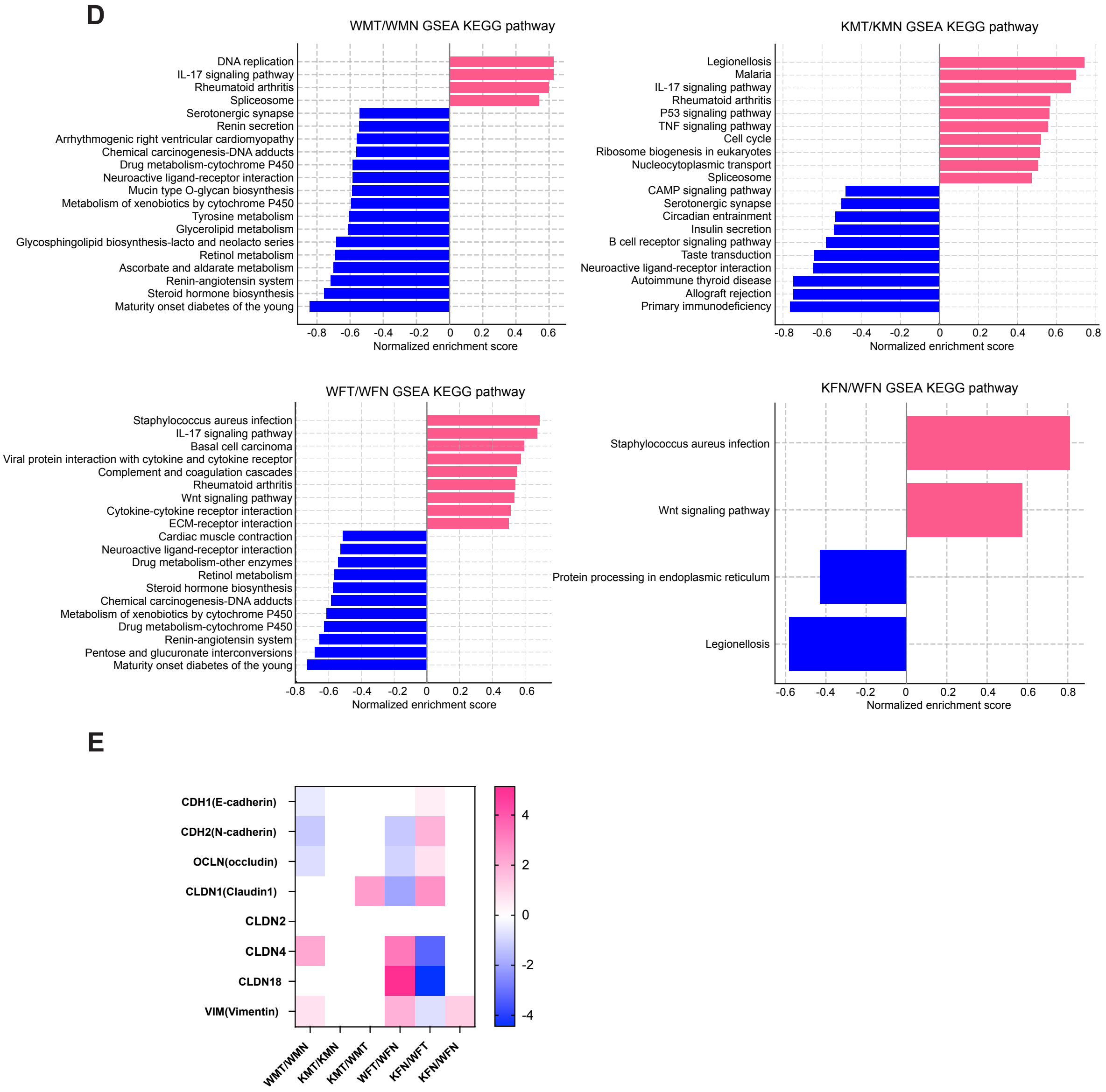


**Fig. S3-2. D,** Gene Set Enrichment Analysis (GSEA) comparing WMT/WMN, KMT/KMN, WFT/WFN, and KFN/WFN pathways using KEGG database categories. The pathways enriched in the dataset are displayed with their corresponding normalized enrichment scores. Pink bars represent pathways that are positively enriched, indicating higher activity or expression in the context of the study, while blue bars represent pathways with negative enrichment, indicating reduced activity or expression. **E,** Heatmap shows the expression levels of various epithelial and mesenchymal markers across different experimental conditions. The markers include E-cadherin (CDH1), N-cadherin (CDH2), occludin (OCLN), and several claudins (CLDN1, CLDN2, CLDN4, CLDN18), along with vimentin (VIM). The color scale indicates expression intensity, with pink showing increased expression and blue indicating decreased expression relative to control levels. Each column represents a different experimental condition, providing insights into the epithelial-to-mesenchymal transition (EMT) status under various treatments or genetic modifications.

Fig.S5.

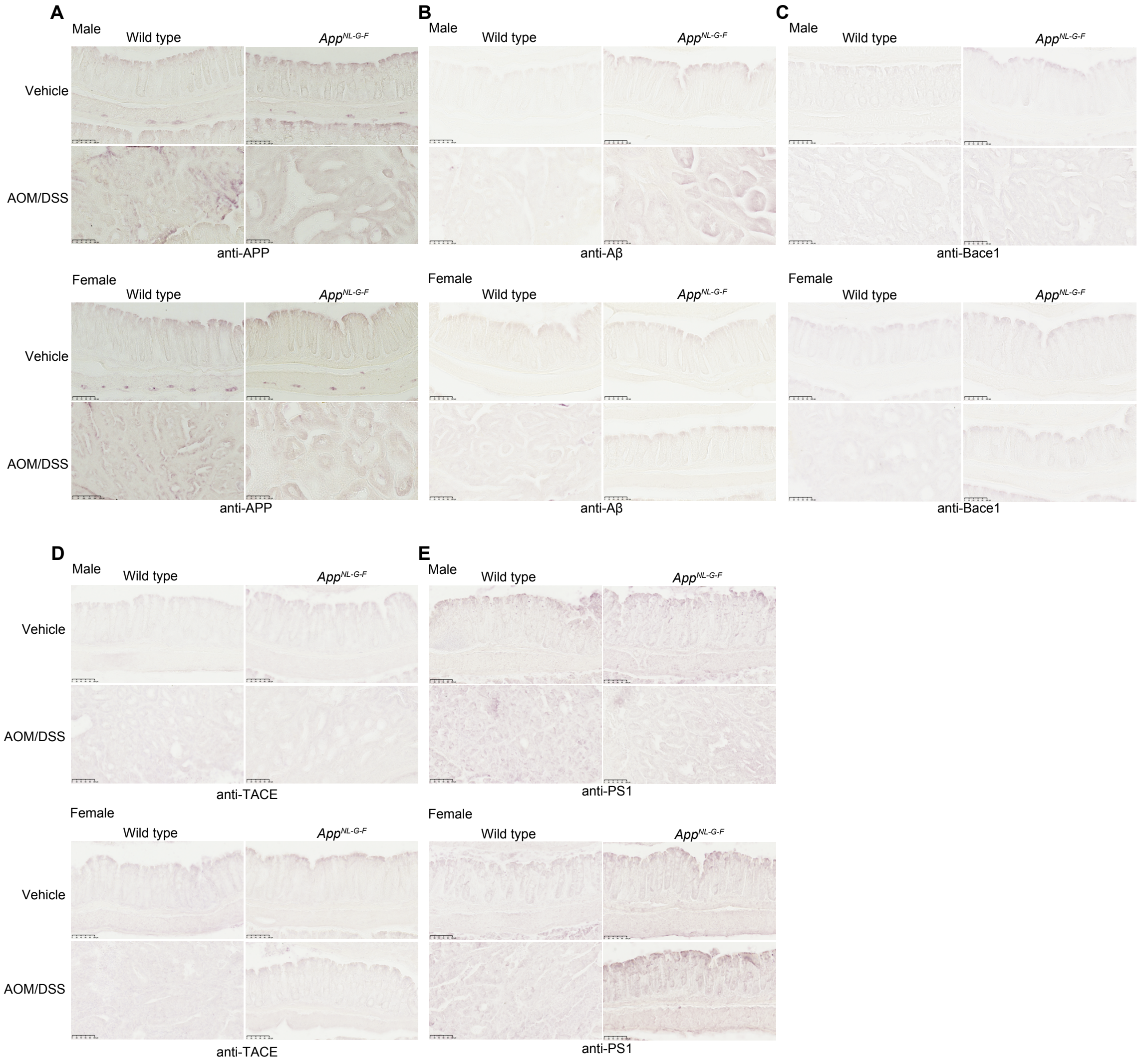


**Fig. S5. Immunohistochemical profiling of AD related proteins in the AOM/DSS colorectal cancer model. A - E,** Immunohistochemical analysis of APP (**A**), Aβ (**B**), BACE1 (**C**), TACE (**D**), and PS1 (**E**) in the colon tissues of male and female wild type and *App*$^{NL-G-F}$ mice treated with vehicle or AOM/DSS. In AOM/DSS-treated panels, images are from cancerous tissues except in *App*$^{NL-G-F}$ female mice. Representative images are shown with 20X magnification.

Fig.S6.

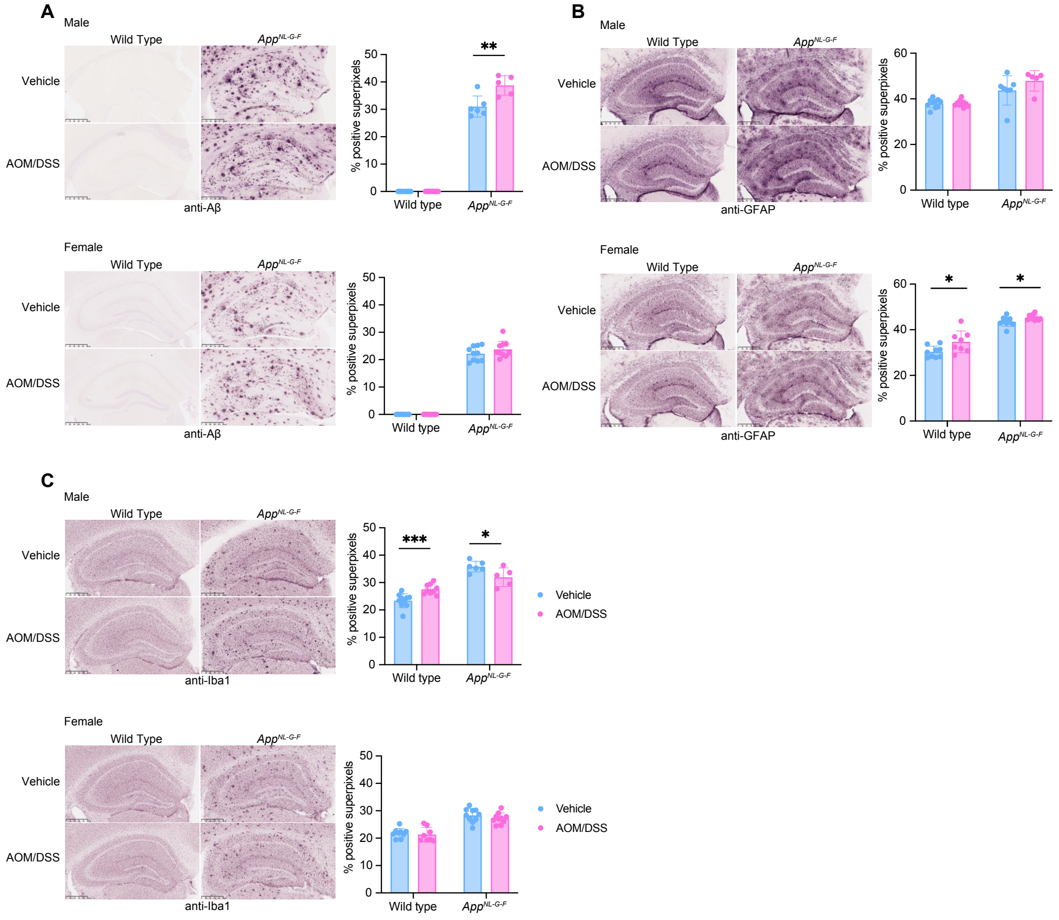


**Fig. S6. Immunohistochemical analysis of gliosis and Aβ in the AOM/DSS mouse model. A - C,** Immunohistochemical analysis of Aβ (**A**), GFAP (**B**), and Iba1 (**C**) in the brains of male and female wild type and *App*$^{NL\text{-}G\text{-}F}$ mice, treated with either vehicle or AOM/DSS. Quantification of positive staining shown in adjacent bar graphs for both genders. Representative images are shown with 4X magnification. 3 different sections of hippocampus/mouse/condition were measured. *p<0.05, ** p<0.01, ***p<0.001, (mean ± SEM, male n=5-10, female n=8-10 replicates)
